# Spatio-temporal dynamics of intra-host variability in SARS-CoV-2 genomes

**DOI:** 10.1101/2020.12.09.417519

**Authors:** Ankit K. Pathak, Gyan Prakash Mishra, Bharathram Uppili, Safal Walia, Saman Fatihi, Tahseen Abbas, Sofia Banu, Arup Ghosh, Amol Kanampalliwar, Atimukta Jha, Sana Fatima, Shifu Aggarwal, Mahesh Shanker Dhar, Robin Marwal, V. S. Radhakrishnan, Kalaiarasan Ponnusamy, Sandhya Kabra, Partha Rakshit, Rahul C. Bhoyar, Abhinav Jain, Mohit Kumar Divakar, Mohamed Imran, Mohammed Faruq, Divya Tej Sowpati, Lipi Thukral, Sunil K. Raghav, Mitali Mukerji

**Author notes:** First author (Ankit K. Pathak, Gyan Prakash Mishra).

## Abstract

During the course of the COVID-19 pandemic, large-scale genome sequencing of SARS-CoV-2 has been useful in tracking its spread and in identifying Variants Of Concern (VOC). Besides, viral and host factors could contribute to variability within a host that can be captured in next-generation sequencing reads as intra-host Single Nucleotide Variations (iSNVs). Analysing 1, 347 samples collected till June 2020, we recorded 18, 146 iSNV sites throughout the SARS-CoV-2 genome. Both, mutations in RdRp as well as APOBEC and ADAR mediated RNA editing seem to contribute to the differential prevalence of iSNVs in hosts. Noteworthy, 41% of all unique iSNVs were reported as SNVs by 30th September 2020 in samples submitted to GISAID, which increased to ∼80% by 30th June 2021. Following this, analysis of another set of 1, 798 samples sequenced in India between November 2020 and May 2021 revealed that majority of the Delta (B.1.617.2) and Kappa (B.1.617.1) variations appeared as iSNVs before getting fixed in the population. We also observe hyper-editing events at functionally critical residues in Spike protein that could alter the antigenicity and may contribute to immune escape. Thus, tracking and functional annotation of iSNVs in ongoing genome surveillance programs could be important for early identification of potential variants of concern and actionable interventions.

**GRAPHICAL ABSTRACT:** 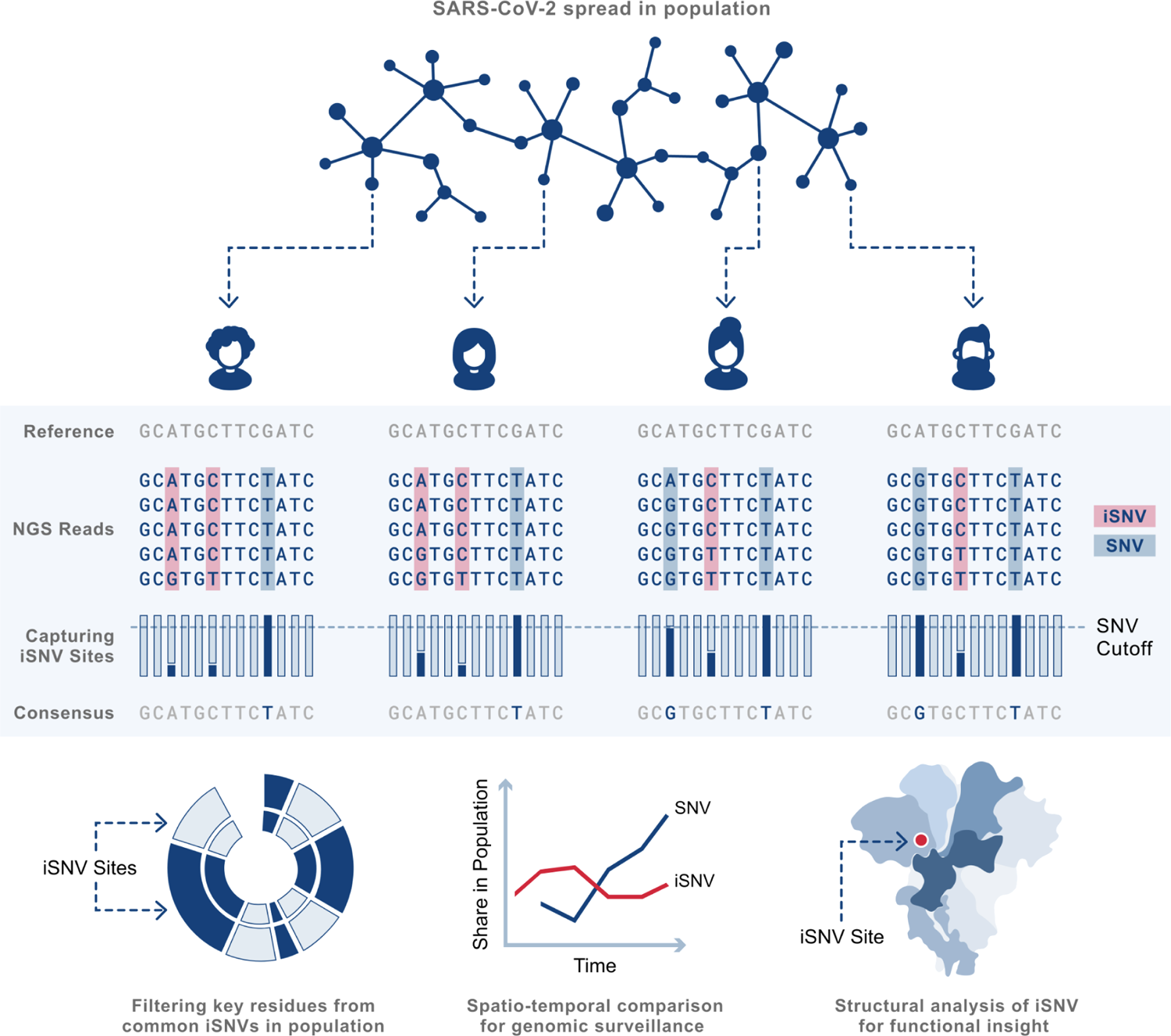

## INTRODUCTION

The SARS-CoV-2 pandemic has inflicted an enormous toll on human lives and livelihoods. As of June 30, 2021, 3.9 million deaths and more than 181 million confirmed cases of COVID-19 have been reported to the WHO from around the globe (1). As the virus accumulates novel mutations, probing millions of viral genome sequences has helped track the continual evolution of SARS-CoV-2. By rigorous cataloging of mutations, genomic surveillance has assisted in predicting the upcoming trajectories of infection by aiding epidemiological models with metrics of transmissibility and virulence of travelling viral strains in a population.

One of the differentiating features of RNA viruses is their exceptionally high nucleotide mutation rates that results in a set of closely related but non-identical genotypes, termed as *quasispecies*, in a host (2–7). Although random mutations have been shown to be largely deleterious (8, 9), some strains may develop enhanced survival ability by evolving mechanisms to resist antiviral responses of the host immune system or drug resistance (10, 11). The fitness of the intra-host viral population has been shown to be dependent on the cumulative contribution of all strains, including haplotypes harbouring minor variants (12). Various factors confer viral heterogeneity within a host like low fidelity of RNA-dependent RNA polymerase (RdRp) and RNA editing by host deaminases combined with the intrinsic high-rates of replication of RNA. Moreover, recent studies have shown mutations in the RdRp to affect the overall genome variability in SARS-CoV-2 (13, 14) whereas RNA editing by host deaminases has been proposed as a major driver of intra-host heterogeneity in the SARS-CoV-2 transcriptomes (15). Understanding these intra-host variations is important as they could potentially alter function through changes in RNA secondary structure, regulatory domains, as well as protein function and thereby govern host-pathogen interactions.

Genomic heterogeneity of the virus within a host can be analysed by capturing intra-host Single Nucleotide Variations (iSNVs), observed as heteroplasmic sites (often referred to in the mitochondrial genome context) in sequence reads. In this study, we accessed samples of COVID-19 diagnosed patients sequenced on an Illumina platform from two different time-periods of the pandemic. In Phase 1, we analysed 1, 347 samples collected latest by June 2020 from China, Germany, Malaysia, United Kingdom, United States and different subpopulations of India to perceive a genome-wide iSNV map in SARS-CoV-2 infected populations. We observed 18, 146 iSNV sites spanning the viral genome, including residues that defined the B.1 and B.6 lineages that were prominent in the listed populations before June 2020. Potential APOBEC and ADAR editing signatures were present throughout the SARS-CoV-2 genome with some samples showcasing ADAR hyperactivity. Further, mutations in RdRp and host-mediated RNA editing showed influence on the differential prevalence of iSNVs between hosts. Structural analysis of iSNVs in the Spike protein revealed a high density of alterations in functionally critical residues that could alter the antigenicity and may contribute to immune escape. Interestingly, 41% of all unique iSNVs identified in these samples were found to be reported as an SNV by 30th September 2020 in one or more samples submitted in GISAID, increasing to ∼80% by 30th June 2021. The likelihood that iSNVs can overtime manifest into SNVs in populations was further substantiated in Phase 2 by analyzing 1, 798 samples sequenced in India between November 2020 and May 2021. In these samples, iSNVs could be recorded to be present in the population in most of the Delta (B.1.617.2) and Kappa (B.1.617.1) lineage-defining genomic positions prior to their fixation as SNVs by February 2021. These results highlight the importance of recording iSNVs as an extension to the genomic surveillance programs to enable more accurate models for viral epidemiology.

## MATERIALS AND METHODS

### Datasets

We processed Illumina next-generation sequencing samples from two different time-periods to understand the magnitude of prevalence of SARS-CoV-2 intra-host Single Nucleotide Variations (iSNVs) in infected individuals across various populations and to explore the possibility of iSNVs getting fixed in a population as a variant. For Phase 1, we accessed 1, 347 publicly available samples submitted in the National Center for Biotechnology Information Sequence Read Archive (NCBI SRA) (16) till 23rd May 2020 from China, USA, Germany, Malaysia and the United Kingdom. Samples from laboratories in Bhubaneswar, Delhi and Hyderabad, that had been sequenced latest by 11th June 2020, were also included to represent SARS-CoV-2 infected East, North and South Indian populations respectively (Table 1). For Phase 2, we analyzed 1, 798 samples sequenced in India between November 2020 and May 2021 for spatio-temporal review of the iSNV states of Delta (B.1.617.2) and Kappa (B.1.617.1) lineage-defining variations.

**Table 1.**
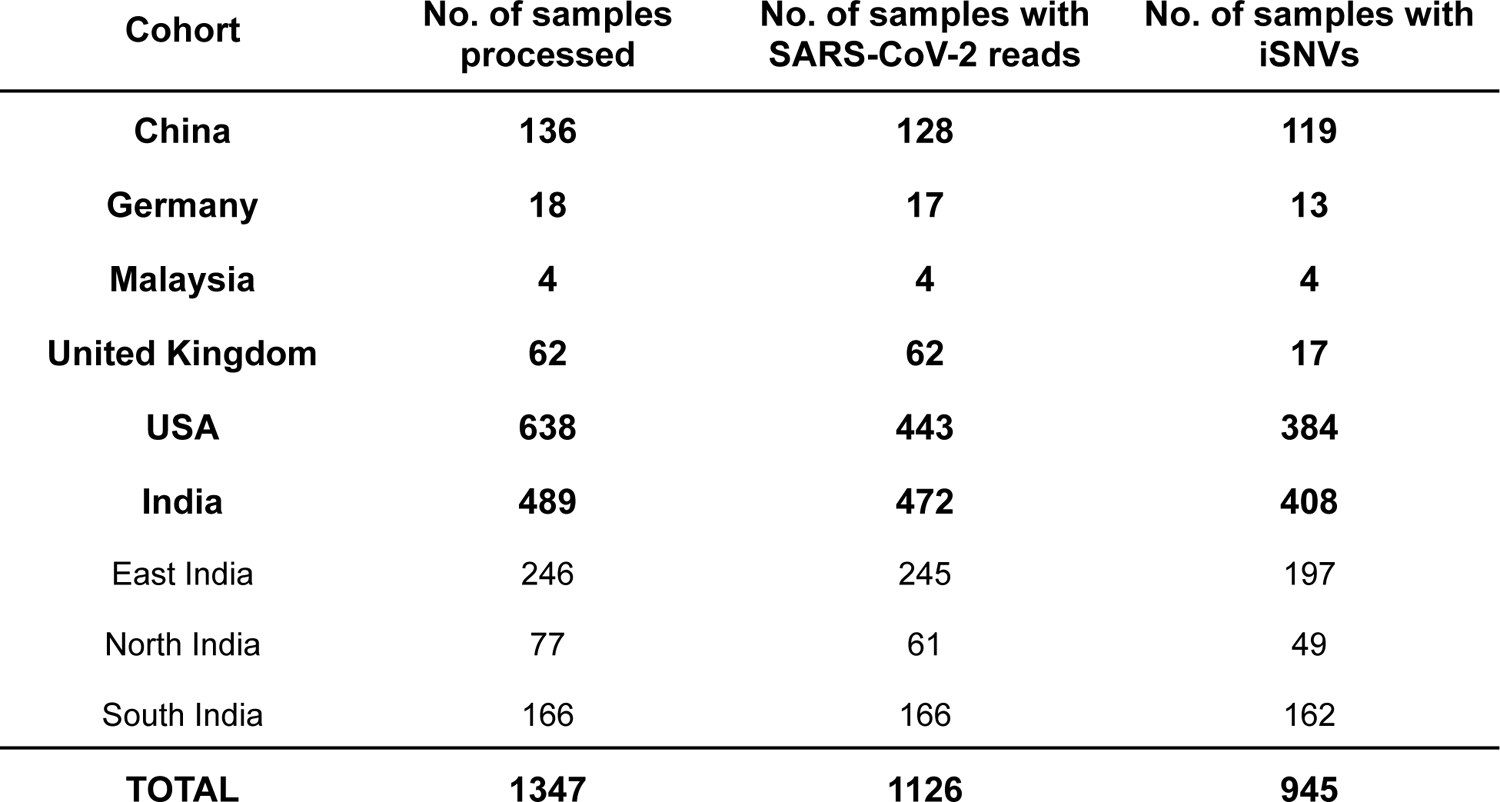
Cohort-wise share of samples for Phase 1. The number of processed samples for each population curated from samples submitted in NCBI SRA till 23rd May 2020 (China, Germany, Malaysia, United Kingdom, USA) and samples sequenced in laboratories from Bhubaneswar, Delhi, and Hyderabad (India) latest by 11th June 2020.

| Cohort | No. of samples processed | No. of samples with SARS-CoV-2 reads | No. of samples with iSNVs |
| --- | --- | --- | --- |
| China | 136 | 128 | 119 |
| Germany | 18 | 17 | 13 |
| Malaysia | 4 | 4 | 4 |
| United Kingdom | 62 | 62 | 17 |
| USA | 638 | 443 | 384 |
| India | 489 | 472 | 408 |
| East India | 246 | 245 | 197 |
| North India | 77 | 61 | 49 |
| South India | 166 | 166 | 162 |
| **TOTAL** | **1347** | **1126** | **945** |

The GRCh37 human reference genome and NC_045512.2 SARS-CoV-2 Wuhan-Hu-1 reference genome were used for all analyses. High-coverage FASTA sequences submitted in GISAID (17) were accessed to get a position-wise share of SNVs in the global SARS-CoV-2 population using the 2019nCoVR database (18).

### Data processing

The sequence files that were retrieved from NCBI SRA were downloaded in the SRA compressed file format and converted to FASTQ format files using NCBI’s SRA-Tools (19). Adapter sequences were trimmed and low-quality reads were filtered using Trimmomatic (v=0.39) for downstream analysis (20). The trimmed reads were then aligned against the human reference genome (GRCh37/hg19) using HISAT2 (21). All paired-end reads unmapped to the human reference genome were filtered out using SAMtools (22) and mapped to the SARS-CoV-2 reference genome (NC_045512.2) using Burrows-Wheeler Aligner (BWA-MEM) (23). Since iSNVs cater to small fractions of reads, to avoid any bias induced by PCR amplification (24), all duplicate reads were removed from the SARS-CoV-2 aligned files using Picard Tools (25). BCFtools (22) was then used to generate the consensus SARS-CoV-2 FASTA sequence. Sample quality scores and alignment statistics were obtained using FastQC (26), QualiMap (27) and SAMtools (22) at each step.

We used REDItools2 (28) to call iSNVs in successfully aligned SARS-CoV-2 reads with a quality score ≥30. Additional position-level parameters in REDItools2 were used to discard some positions on the basis of site quality. The sites whose reads did not achieve a mean quality score of ≥33 were rejected and sites that were located in long homopolymeric regions of length ≥4bp (Supplementary Dataset S1) were excluded due to known sequencing errors in these regions (29). From the REDItools2 output, high-quality iSNV events were further filtered out using the following thresholds: number of minor allele reads ≥5, base coverage ≥20 and minor allele frequency ≥0.005. In order to test the pipeline’s efficiency in identifying iSNVs in SARS-CoV-2 genome, we additionally downloaded and processed 500 single-end NGS samples sequenced in replicates on an Illumina platform from NCBI SRA with the Project ID PRJNA655577. We observed a high concordance between the replicates (Pearson’s r=0.989, Supplementary Figure S1) in sites qualifying the pipeline thresholds. Thus, we used these cutoffs as a qualifying criterion for defining a heteroplasmic site as an iSNV in our subsequent analyses.

### SARS-CoV-2 virus culture in Vero E6 cells

#### Virus isolation

Virus isolation from patient’s swab samples was done following Harcourt. et al, 2020 (30). Vero E6 cells were grown in a 6-well plate at 80% confluency in DMEM supplemented with antibiotics (Penicillin-Streptomycin) and 10% Fetal bovine serum. Next day, Vero E6 cells were infected with 11 SARS-CoV-2 isolates at MOI of 1 in serum free media for 1 hour at 37°C with gentle rocking every 10-15 mins. One hour post infection, the inoculum was aspirated and cells were washed with PBS and supplemented with fresh complete media. The culture supernatant was aspirated 48 hours post infection, the cells were washed with 1x PBS and collected in Trizol reagent for total RNA extraction. SARS-CoV-2 virus isolation and culture was conducted in a biosafety Level-3 facility according to the biosafety guidelines issued by the Department of Biotechnology, Government of India.

### qRT-PCR for viral load estimation and mRNA expression of APOBEC3B and ADARB1

RNA isolation from culture cells was performed using TRIzol Reagent (Invitrogen, cat. no. 10296010). The isolated RNA was subjected to qRT-PCR for determining the viral load. We performed one-step multiplex real-time PCR using TaqPathTM 1-Step Multiplex Master Mix (Thermo Fisher Scientific, cat. no. A28526), targeting three different SARS-CoV-2 genes with primer and probe sets specific for Spike (S), Nucleocapsid (N), and Open Reading Frame 1 (ORF1). To check the expression of APOBEC3B and ADARB1, RNA from wild-type (mock) and Vero E6 cells infected with 2 patient virus isolates was converted to cDNA using reverse transcriptase from qiaseq SARS-CoV-2 primer pool (Qiagen kit, cat no. 333896). APOBEC3b (sense: 5’-TCCACAGATCAGAAATCCGA-3’; antisense: 5’-CCATAGAGGATGGGTTCGT-3’) and ADARB1 (sense: 5’-AAACATGAGTTCCAGCAGC-3’; antisense: 5’-CCCATTGGAGAGCTGAGAG-3’) primer sequence were used to estimate the mRNA expression. 18S rRNA Primers (sense: 5’-CTACCACATCCAAGGAAGCA-3’ and antisense: 5’-TTTTTCGTCACTACCTCCCCG-3’) were used as internal control. qRT-PCR was performed using the Applied Biosystems (QuantStudio 6) machine.

### Viral RNA library preparation and sequencing

Amplicon libraries for viral genome sequencing were prepared using QIAseq FX DNA Library Kit and QIAseq SARS-CoV-2 Primer Panel (Qiagen, cat. no. 180475, cat. no. 333896) as instructed by the manufacturer’s manual, and the library was subsequently sequenced using Illumina NextSeq 550 platform. The adapter sequence used for each sample was compatible with Illumina sequencing instruments with 96-sample configurations (Qiaseq unique dual Y-adapter kit). The average insert length was in the 250–500 bp range. To characterize iSNVs, the *Chlorocebus sabaeus* genome (chlSab2 assembly) was used inplace of the human genome in the pipeline.

### Identification of samples with potential hyper-editing events

Z-score values based on the distribution of number of iSNVs per sample were calculated for each population using the Python package SciPy (31). Samples with a Z-score value > 3 were classified as hyper-edited in each cohort (Figure 3A).

In order to categorize the specific amino acid change and the proteins containing the iSNVs, they were annotated using SnpEff version 4.5 (32). Conservation analysis of the full-length sequences of proteins was done on the basis of the six other coronaviruses (HCoV-229E, NL63, OC43, HKU1, MERS-CoV, SARS-CoV). The multiple sequence alignment of seven protein sequences was performed by clustal-omega (33). Conservation calculation of Spike amino acids was done using the ConSurf tool (34). The high-risk hyper-edited sites resulting in a single amino acid change in Spike proteins were screened and prioritized on the basis of sequence conservation and frequency i.e. number of times it is present in the total 25 hyper-edited samples. ConSurf conservation scores above 6 were considered and a total of 192 variants were filtered for mapping onto the structure to understand the effect on protein function and structure. We utilized PyMol (35) to determine interface residues from the available crystal structures. PDB 6M0J (36) and 6M17 (37) for ACE2 RBD binding and PDB 6W41 (38) and 7BZ5 (39) for antibody binding residues were used from Research Collaboratory for Structural Bioinformatics Protein Data Bank (RCSB PDB) (40).

### Data handling and visualizations

Custom python scripts were used to automate the process of downloading and retrieving samples from NCBI SRA as well as to process samples through the pipeline. Python libraries such as NumPy (41), Pandas (42), Matplotlib (43) and Seaborn (44) were used for data handling and visualizations. SciPy (31) was used to compute pairwise two-sided t-tests and to calculate distribution statistics and Pearson’s correlation coefficients.

## RESULTS

### Intra-host Single Nucleotide Variations (iSNVs) in SARS-CoV-2 genomes

We analysed 1, 347 transcriptomic samples obtained from patients diagnosed with COVID-19 by June 2020 from different populations (Table 1). 1, 126 samples could be aligned to the SARS-CoV-2 reference genome. To ensure specificity of iSNV detection, the filtering criteria and cutoffs used in the sequence reads were established by analysing an additional set of 500 samples sequenced in replicates (Supplementary Figure S1). 945 of the 1, 126 samples (∼84%) harboured one or more iSNVs with frequencies ranging between 0.005 and 0.80 (Supplementary Dataset 3a, 3b). We recorded a total of 79, 068 iSNVs with a median of ∼19 iSNVs per sample (Supplementary Figure S2A), revealing extensive heteroplasmy in samples. We did not observe a significant correlation between the mean sample coverage depth and the number of iSNVs per sample suggesting that these events are unlikely to be a consequence of sequencing artefacts (Pearson’s r=0.277, Supplementary Figure S2C). Further, in a sample subset, we did not find any correlation between the number of iSNVs in a sample and the Ct values for gene N (Pearson’s r=-0.173, *p*=0.065), ORF1 (Pearson’s r=-0.153, *p*=0.105), and S (Pearson’s r=-0.132, *p*=0.162) (Supplementary Figure S2G).

Nucleotide changes due to iSNVs presented a skewed pattern, with A-to-G transition accounting for about a quarter of all changes (Figure 1A). Moreover, A-to-G changes were observed in disproportionate numbers in few samples while C-to-T changes were more consistently observed with a median of 5 events per sample. iSNVs contributed to variability throughout the SARS-CoV-2 genome, both in the coding as well as in the non-coding regions. A total of 23, 392 unique iSNVs were recorded at 18, 146 genomic positions (4, 909 polymorphic) (Supplementary Dataset S4) in one or more samples distributed almost evenly in all genes and protein-coding domains of the genome (Supplementary Figure S3A). Though the frequency spectrum of iSNVs was nearly uniform with respect to the nature of amino acid sequence change, a large fraction of the sites lead to non-synonymous variations (Supplementary Figure S2E). Gain of stop codons that could alter the amount of functional proteins from the viral genomes were also observed.

**Figure 1.**
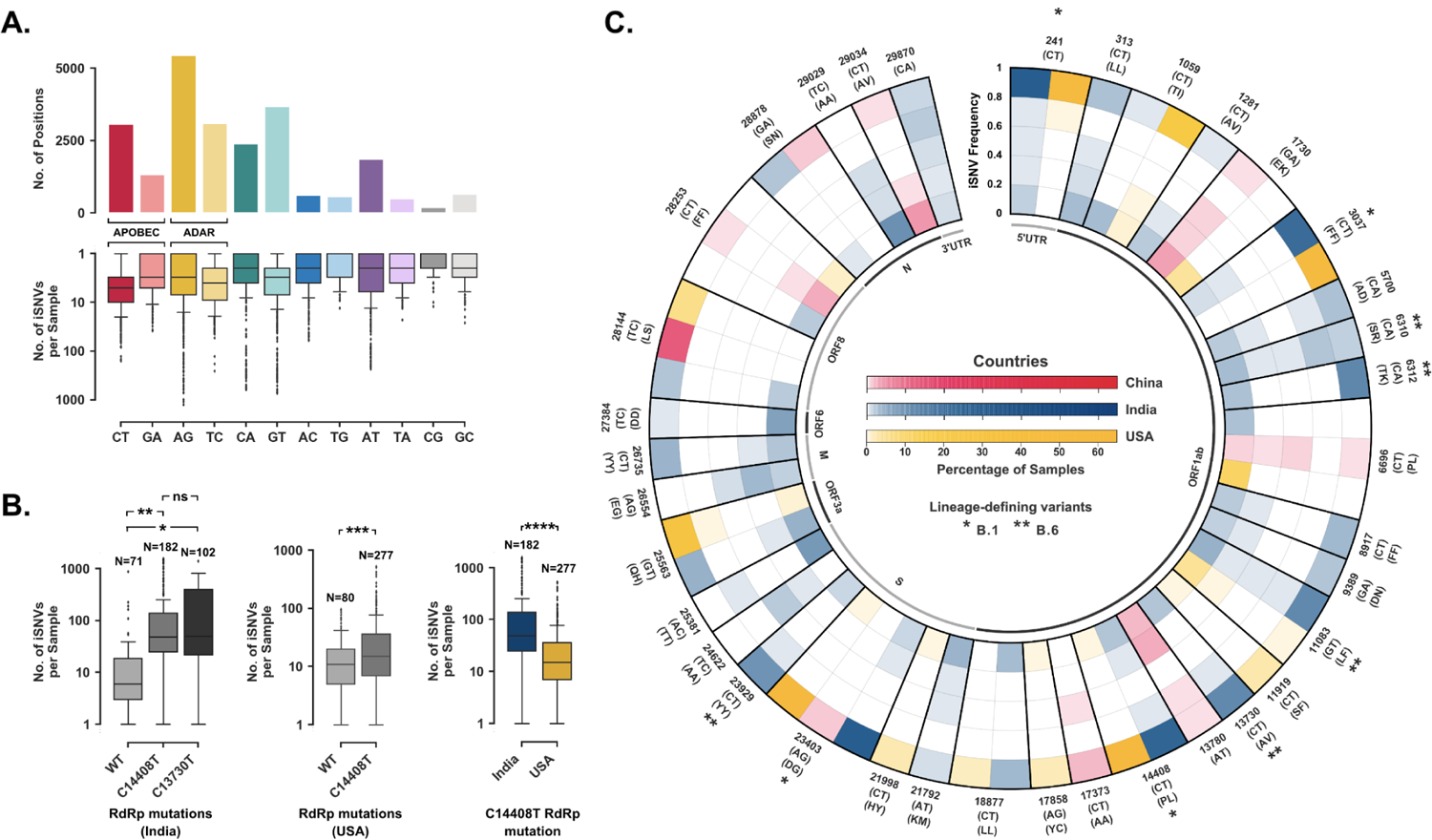
Spectrum of iSNVs in Phase 1 samples. (**A**) Split plot depicting the distribution of unique iSNV sites (n=23392) with respect to the nucleotide change in the SARS-CoV-2 genome and across samples (**B**) Box plots showing comparison of iSNV prevalence between wild-type and RdRp mutant samples in India and USA (* *p* < 10^-6^, ** *p* < 10^-3^, *** *p* < 0.05, **** *p* < 10^-10^). (**C**) Radial plot showcasing frequency distribution for selected iSNV sites in China, India and USA denoted in Red, Blue and Yellow respectively. Each concentric ring depicts an iSNV frequency range of 0.2 and the different colour gradients depict different populations with percentage of samples at a given position in each bin. The outer labels denote the position of change, nucleotide change and amino acid change. Variations that define the B.1 and B.6 lineages have been marked (*) and (**) respectively.

### Putative role of host and viral factor in iSNV incidence

Various factors have been shown to confer intra-host variability in many families of RNA viruses. Low-fidelity of RNA-dependent RNA polymerase (RdRp) has been established as a major contributor to this heterogeneity. Recent reports showed that mutations in the RdRp of SARS-CoV-2 may impact the number of mutations in the viral genome (13, 14). We observed significant difference in the number of iSNVs between the wild-type and C14408T mutant RdRp samples in both India (*p* < 10^-3^) and USA (*p* < 0.05) with higher prevalence in the mutant RdRp (Figure 1B).

A bias towards the C-to-T/G-to-A and A-to-G/T-to-C nucleotide changes suggests RNA editing activity in both the strands of SARS-CoV-2 genome by the APOBEC (apolipoprotein B mRNA editing catalytic polypeptide-like) and ADAR (Adenosine Deaminase RNA Specific) family of enzymes respectively (Figure 1A). APOBECs target single-stranded nucleic acids for deamination of cytosines into uracils (C-to-U) reflecting as a C-to-T transition in the reference genome. ADARs, on the other hand, deaminate adenines into inosines (A-to-I) on double-stranded RNA (dsRNA). Inosine preferentially base pairs with cytidine which leads to incorporation of guanine in subsequent replication and an A-to-G transition. Editing could also occur in the negative strand of SARS-CoV-2 genome that would reflect as G-to-A (APOBEC) and T-to-C (ADAR) changes in the reference genome. Both C-to-T/G-to-A and A-to-G/T-to-C combinations were present throughout the genome (Supplementary Figure S3B) and contributed significantly to synonymous and missense changes, whereas only the former contributed to stop gains (Supplementary Figure S2F). Within the C14408T RdRp mutation carrying samples in India and USA, we observed differential prevalence of iSNVs (*p* < 10^-10^) (Figure 1B). We also performed qRT-PCR analysis of APOBEC3B and ADARB1 expression in Vero cells where we clearly observed high and constitutive ADARB1 expression in both mock as well as in SARS-CoV-2 infected Vero cells (n=2), although APOBEC3B expression could not be captured (Supplementary Dataset S5). Further, we infected Vero cells with independent isolates of SARS-CoV-2 from 11 samples in the East India cohort. Comparing iSNVs at the 0th-passage and 7th-passage, we found significant accumulation of iSNVs in just 7 passages (Supplementary Dataset S6). Interestingly, the relative pattern of iSNVs with respect to the nucleotide change remained similar (Supplementary Figure S6).

### Contribution of iSNVs to the spectrum of SARS-CoV-2 diversity across populations

To understand the contribution of iSNVs in the diversity of SARS-CoV-2 across various populations, we plotted a radial heatmap of the top 1% frequently observed genomic positions exhibiting a wide range of iSNV frequencies in major cohorts (Figure 1C). Each concentric ring represents an iSNV frequency range of 0.2 and the different colour gradients depict various populations with percentage of samples at a given position in each bin. Thus, the innermost ring represents a cohort-wise share of samples which show variations in a small fraction of reads (≤0.2) at a given position whereas the outermost ring represents the ones where the variant seems to have achieved near fixation (>0.8) and would be reported as an SNV in the population. Most of these positions were observed to be shared between global populations and also within Indian subpopulations (Supplementary Figure S4). Interestingly, many samples carried iSNVs at positions which defined the B.1 lineage (C241T, C3037T, C14408T and A23403G) and the India specific B.6 lineage (C6310A, C6312A, G11083T, C13730T and C23929T) (Figure 2A) (45, 46). Noteworthy, all B.1 lineage-defining variants depicted a C-to-T or an A-to-G nucleotide change.

**Figure 2.**
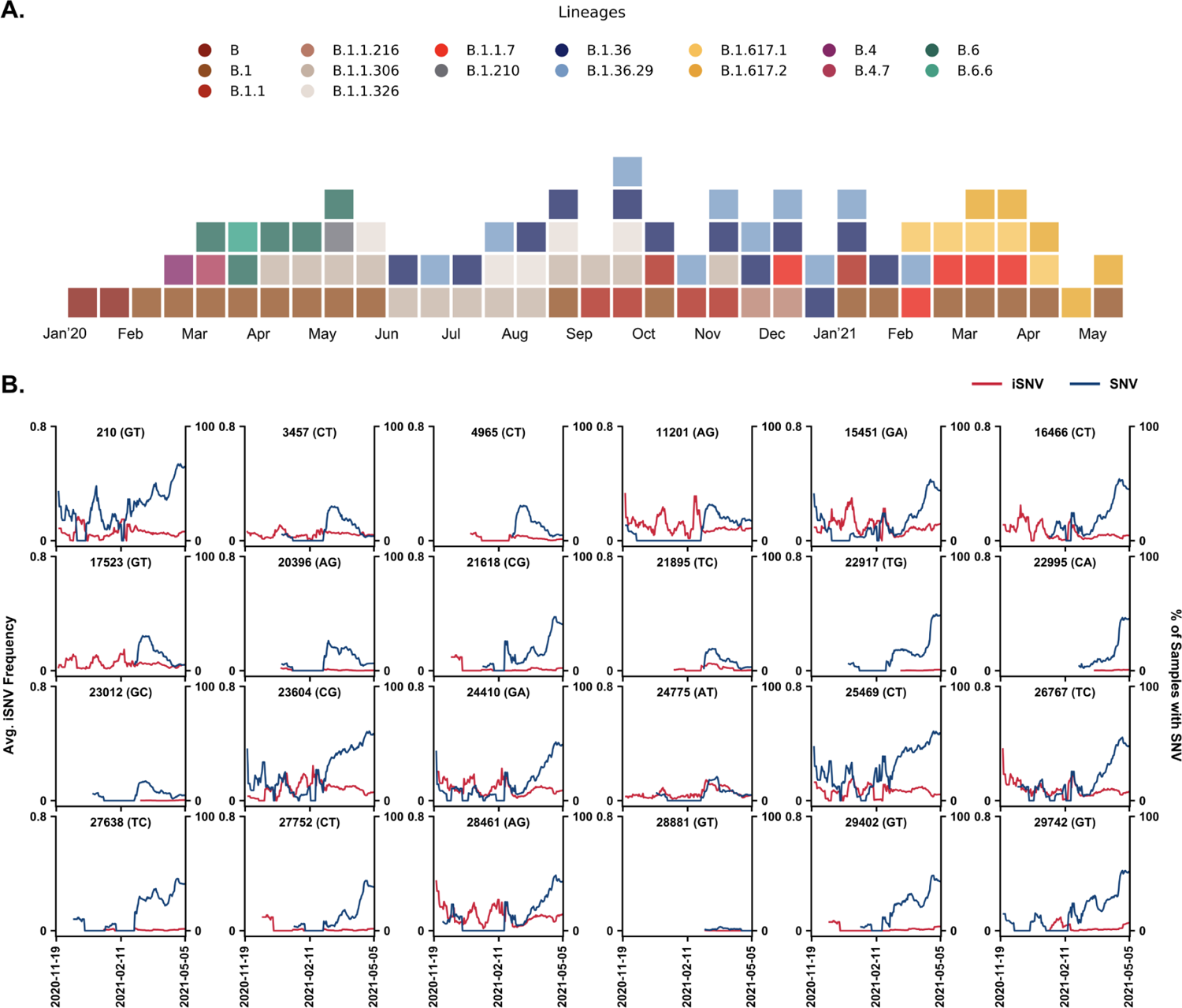
Spatio-temporal dynamics of iSNVs. (**A**) Temporal (fortnightly) distribution of prominent lineages identified in SARS-CoV-2 samples submitted pan-India till May 2021. Only lineages with at least 10% representation in the total samples submitted in a given fortnight period were selected. (**B**) Line plot showing temporal trends of iSNV frequency and incidence of SNV for Delta and Kappa lineage-defining positions in India between November 2020 to May 2021. The left y-axis denotes the average iSNV frequency and the right y-axis denotes the percentage of samples with SNVs on a 14-day rolling basis. Red and blue lines illustrate the average iSNV frequency and the percentage of samples with SNVs at the site respectively.

### Spatio-temporal dynamics of iSNVs

Though most of the sites seem to convert to a fixed variant in one or all populations (Figure 1C, Supplementary Figure S4), positions like 1707, 15435, 24622, 26554 and 29029 that depict an increasing trend in the iSNV frequency in few cohorts suggest that some iSNVs could get fixed in the population at a later date. We found significant overlap of iSNV sites with SNVs that have been reported in sequences submitted in GISAID. 41% of the 23, 392 unique iSNVs identified in the samples were recorded as SNVs in one or more samples submitted in GISAID by 30th September 2020 with the exact change in the nucleotide sequence, increasing to ∼80% by 30th June 2021 (Supplementary Dataset S4). As of June 2021, WHO had listed 4 lineages as Variants of Concern (VOC) and 7 as Variants of Interest (VOI). ∼74% of all positions defining these lineages were recorded as iSNV sites in one or more samples including 75.6% of all Spike mutations (Supplementary Dataset S7). Given the origin and start of the outbreak of Delta (B.1.617.2) and Kappa (B.1.617.1) lineages in February 2021 in India (Figure 2A), we analysed 1, 798 samples sequenced between November 2020 to May 2021 from different parts of the country. We found that at majority of the Delta and Kappa lineage-defining positions, SNVs appeared coinciding with or after the iSNVs first surfaced in the population (Figure 2B). This further strengthens the postulate that iSNVs may introduce SNVs into the travelling viral strains in a population which may beget newer lineages with heightened properties of transmissibility and virulence as seen with the Delta lineage.

### Potential functional consequence of hyper-editing on Spike protein

We observed a few samples in each cohort to be substrates for potential hyper-editing events as reflected by an excessive number of iSNVs. In order to identify these population outliers, we computed the Z-score values based on the distribution of the number of iSNVs per sample (Figure 3A). In these samples, we observed A-to-G substitutions accounting for about one-third of all changed genomic positions implying extensive ADAR mediated editing activity (Figure 3B). Of the 11, 420 annotated variants, 1, 481 correspond to protein-coding variants within the Spike protein.

**Figure 3.**
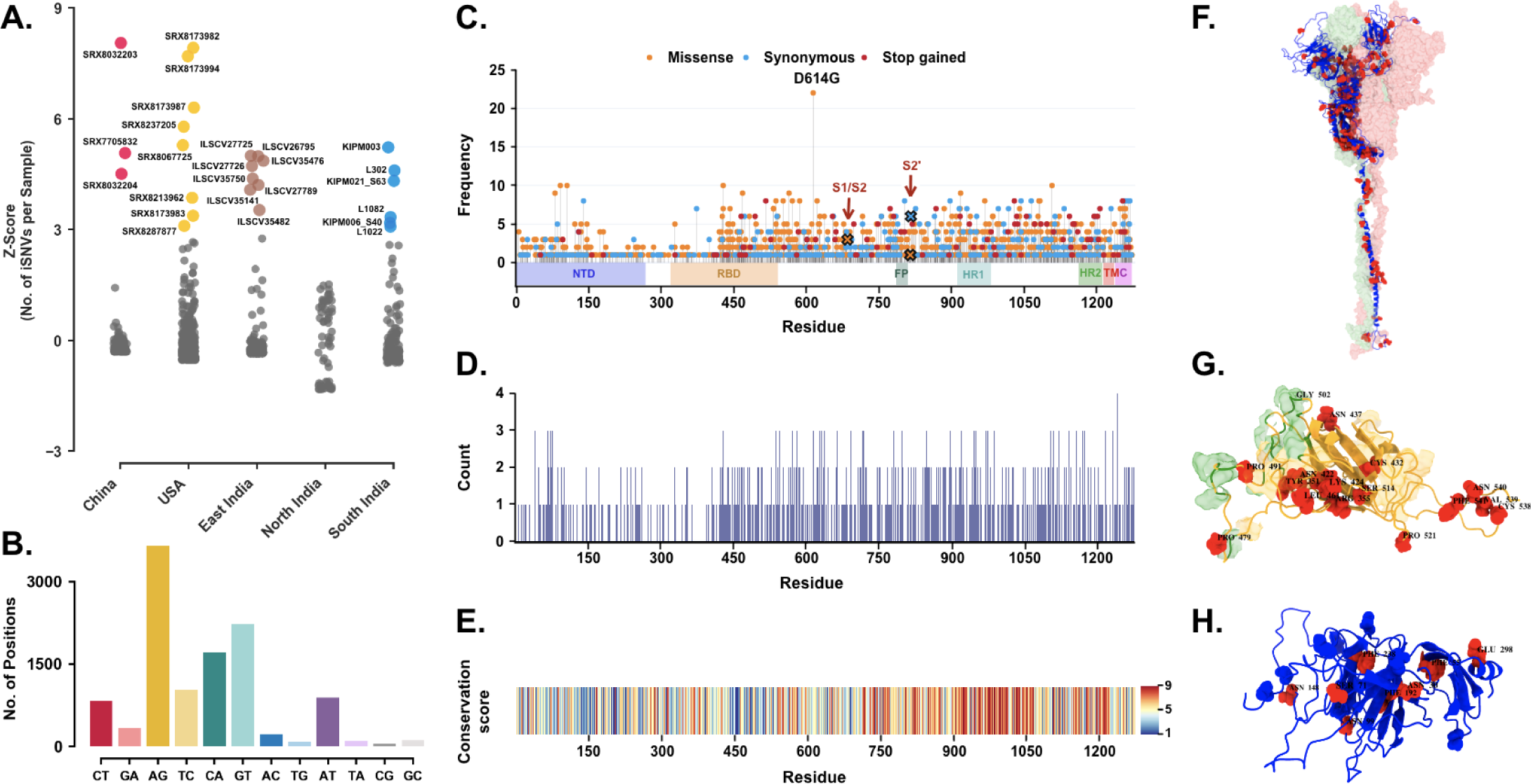
Functional impact of hyper-editing on Spike protein. (**A**) Dot plot illustrating cohort-wise outliers in Z-score values based on the distribution of number of iSNVs per sample (**B**) Bar plot representing distribution of iSNV sites (n=11392) with respect to nucleotide changes in the SARS-CoV-2 genome in hyper-edited samples (**C**) Needle plot depicting the distribution of protein-coding changes due to iSNVs in Spike protein. Non-synonymous, synonymous, and stop variants are shown in orange, blue, and red respectively. The length of the needle depicts the occurrence of altered residues out of the total of 25 hyper-edited samples. Protein domain architecture is indicated as horizontal boxes (**D**) Counts of amino acid substitutions at each residue location in the Spike protein (**E**) Conservation score of residues in the Spike protein. A conservation score of 9 denotes a highly conserved while a score of 1 denotes a highly variable position (**F**) Selected iSNVs with missense changes mapped onto the Spike trimer structure where red dots represent altered sites. The other two chains of the trimer are shown in surface representation (**G**) A close-up view of the RBD domain in surface representation harbouring the altered sites (iSNVs) in hyper-edited samples. Green highlights the ACE2 binding surface and orange highlights the antibody binding surface (**H**) A close-up view of the NTD domain with altered residues shown in red and N-glycosylation sites in blue. Altered residues are labelled in black text: amino acid name and position.<colcnt=2>

To understand how iSNVs could impact function through amino acids substitutions in hyper-edited samples, we examined the mutational spectra of the Spike protein in more detail, given that the initial host-viral response is triggered by the attachment of the Spike protein of SARS-CoV-2 with the host ACE2 receptor, disrupting the host cell membrane and activating viral entry (47). Amongst the 1, 483 iSNVs in the Spike protein, 1, 068 were found to be non-synonymous, 322 synonymous and 91 stop-coding. Figure 3C depicts the frequency of these sites on different functional protein domains, including the critical receptor-binding domain (RBD). The most frequently altered variants were D614, Y91, I105, and D428. The D614G mutation (one of the B.1 lineage defining variants) is near the S1/S2 cleavage site and has already been shown to be more geographically spreading than the D614 type (48).

We looked at other Spike mutations in more detail with respect to their functional consequences. Most striking, some amino acid sites within Spike seem to evolve into more than one type of variant. In Figure 3D, we depict these hyper-variable sites, with residue A879, N1108, S1239, D1260 evolving into either of >4 amino acid substitutions. The conservation of each residue i.e., how variable that site is amongst closely related coronavirus species was also calculated. For instance, the HR1 region and residues near the transmembrane are observed to be more conserved while NTD and RBD domains have highly variable regions, in that, especially the RBM motif (residue 437-508) seems to be less conserved (Figure 3E). Interestingly, RBM was also found to be variable in our study where specific amino acids at 442, 465, 468 conferred variations in ∼11 out of 25 hyper-edited samples. The highly conserved amino acids represent a similar function of the protein across species but the substitution of critical amino acids around functional sites like RBM motif might lead to evolved or newer biological activities.

In a recent study, accelerated evolution of the virus over time leading to a repeated resurgence in an immunocompromised infected patient has been reported (49). Genomic analysis revealed that the individual had not been infected multiple times. Instead, the virus had lingered and rapidly mutated in the body leading to non-synonymous changes predominantly in the Spike region. The Spike protein and RBD harboured 57% and 38% of the total changes despite occupying only 13% and 2% of the virus genome respectively. The changes also included Q493K and F486I that corroborated with an earlier study that had recognised additional mutations (N74K, F79I, T259K, K417E, K444Q, V445A, N450D, Y453F, L455F, E484K, G485D, F490L, F490S, H655Y, R682Q, R685S, V687G, G769E, Q779K and V1128A) to be implicated in antibody resistance (50). Besides, N439K mutation in Spike has also been shown to be involved in immune escape from a panel of neutralizing monoclonal antibodies in another study (51). 20 out of these 23 positions overlap with sites that have been shown to carry iSNVs in our samples (Supplementary Dataset S8).

## DISCUSSION

The course of the current pandemic has seen huge efforts in sequencing SARS-CoV-2 genomes from COVID-19 diagnosed patients and has helped catalogue the spread of mutations since the initial strain reported from Wuhan back in December 2019. Genomic surveillance has become an essential aid to epidemiological models and has facilitated prior predictions of variation(s) of concern through observations on inter-host differences in the viral genome over a period of time. In this study, we explored intra-host variability in the SARS-CoV-2 genome at spatio-temporal scales as an extended tool for genomic surveillance and examined factors contributing to its prevalence and its downstream effects.

Intra-host Single Nucleotide Variations (iSNVs) can alter the fitness of the strains by influencing its virulence, infectivity or transmissibility properties which may be both beneficial or detrimental to the virus and lead to fixation or removal of alleles from the populations. Of the 23, 392 unique iSNVs recorded in samples collected latest by June 2020, 41% and ∼80% had been recorded as an SNV in samples submitted in GISAID (17) till 30th September 2020 and 30th June 2021 respectively (Supplementary Dataset S4). Given the global spread of the infection, even within the relatively small subset of samples analysed in this study from select population groups which were collected in the first few months of the pandemic, ∼74% of all the variants listed by the WHO as VOCs or VOIs by June 2021 (52) had been recorded as an iSNV site in one or more samples (Supplementary Dataset S7). Recent studies have also suggested that the intra-host heterogeneity of SARS-CoV-2 genome in individuals may be transferable (53–55), and given that a minuscule percent of population is responsible for most of the local transmission of SARS-CoV-2 infection (56–58), there arises a possibility of iSNVs harbouring genome to be passed through a super-spreading event. This, in an isolated population, might also lead to an altered haplotype with some unique variations leading to fixation or removal of alleles from the population (59). By temporally capturing iSNVs and the consequential variant appearance in the population, we could determine iSNV sites in the East Indian population which possibly transitioned into an SNV (Supplementary Figure S5). Further analysing samples sequenced in India between November 2020 to May 2021, we found majority of Delta and Kappa lineage-defining positions to carry iSNVs before the incidence of the subsequent SNV in the population (Figure 2B). This indicates that conjoint analysis of iSNVs with such gain and loss of variants in spatio-temporal scales might provide some clues of the evolvability of sites and may prove to be an important asset in genomic surveillance. Moreover, co-occurrence of such events in distinct isolated populations may result in the independent origin of lineages with similar properties.

Although the extent of variability differs amongst individuals, there seem to be preferred iSNV sites that are shared across populations (Figure 1C, Supplementary Figure S4). By the end of 2020, the B.1 lineage had become the most prevalent in most parts of the world with near disappearance of other lineages which were once the predominant strain in certain regions. For example, the B.6 lineage which was reported to be primarily endemic to India (45, 46) in April 2020 had disappeared from the entire population by October 2020 (Figure 2A). Given that all B.1 lineage-defining positions (241, 3037, 14408 and 23403) are iSNV sites with C-to-T and A-to-G changes (Figure 1C), host-mediated RNA editing might have fueled the positive selection of the B.1 lineage. The APOBEC and ADAR family of enzymes are extensively reported to edit different families of viral genomes (60–64) including the coronaviruses (65). SARS-CoV-2 being zoonotic and of recent origin, seems to harbour a large number of sites that could be potentially targeted by the host editing machinery. The signatures of these enzymatic activities seem to be reflected in the evolving SARS-CoV-2 genomes (Figure 1A) which corroborates with an earlier report (15). The RT-qPCR expression data revealed high ADARB1 expression in SARS-CoV-2 infected Vero cells indicating its role in genome mutagenesis (Supplementary Dataset S5). Though we observed potential ADAR hyperactivity in some individuals, APOBEC-mediated C-to-T changes were more consistently observed in our samples (Figure 1A) which may explain the overrepresentation of C-to-T SNVs in the SARS-CoV-2 populations (66, 67). Besides, we also observed substantial abundance of G-to-T and C-to-A substitutions in some samples. These might be due to Reactive Oxygen Species (ROS), as hypothesized in a recent study (68). ROS activity oxidizes guanine to 7, 8-dihydro-8-oxo-2’-deoxyguanosine (oxoguanine) that base pairs with adenine, leading to G-to-T transversions. This change on the negative strand would be reflected as a C-to-A transversion in the reference genome. ROS also has its role implicated in mutagenesis of many other virus families (69).

Low fidelity of RNA-dependent RNA polymerase (RdRp) is known to be a major driver of intra-host heterogeneity of RNA viruses. Correlating the number of iSNVs against mutations in RdRp revealed differential prevalence of iSNVs between the wild-type and C14408T mutant RdRp in samples from both India as well as USA, indicating positive selection of a more error-prone RdRp to maintain an active buffer of minor variants (Figure 1C). The difference in the prevalence of iSNVs could also arise out of varied RNA editing across populations (Figure 1C), highlighting the active role of the host. The editing enzymes have been shown to display genetic variability in populations, perhaps a consequence of selection based on pathogenic loads (70, 71). Transmission of viruses through different host populations could thus shape the mutational landscape and govern the evolution of the viral genomes.

Ongoing selection in the host may contribute to the fixation of some of the sites as variants in SARS-CoV-2 genomes with potential consequences on the functioning of some of the viral proteins. Structural analysis of iSNVs that occur in Spike shows that more than 40% of these variants result in a substantial change in the property of these amino acids (Supplementary Table S1). Changes in specific amino acids, especially mutations close to functional sites, may lead to drastic changes in protein structure as they can either introduce significant change in the sidechain or charge of the residue. Also, it has been shown that APOBEC influenced C-to-T substitutions elevate the frequency of hydrophobic amino acid coding codons in SARS-CoV-2 peptides (72). We also mapped 192 iSNVs onto the protein structure by utilizing a full-length Spike protein model of closed conformation (Figure 3F). Some iSNVs are located on the surface of the receptor-binding domain (RBD) which is responsible for binding to the ACE2 receptor in the host cell. These would directly impact the binding of Spike protein to the receptor and might have a putative effect on the binding interface residues shown in surface representation (Figure 3G). Noteworthy, we observed iSNVs in ∼87% of the sites in the Spike protein that have been recently reported to confer antibody resistance (Supplementary Dataset S8). These mutations can have major implications in vaccine response as they could alter the immunogenicity of the antigenic RBD peptide leading to differential antibody titers in infected individuals (73). Overall, our findings indicate a comprehensive survey for one of the SARS-CoV-2 proteins and we propose that some of these distinct patterns of sites in hyper-edited samples may directly interfere with specific functional output.

In conclusion, temporally tracking within-host variability of the virus in individuals and populations might provide important leads to the sites that are favourable or deleterious for virus survival. This information would be of enormous utility for predicting the spread and infectivity of viral strains in the population. Conjoint analysis with the intra-host variability of the SARS-CoV-2 genome should be the next step.

### DATA AND CODE AVAILABILITY

The commands used in the pipeline, automated codes, detailed individual sample information and statistics are available at the following GitHub repository — https://github.com/pxthxk/iCoV19.

## FUNDING

This work was funded and supported by the Council of Scientific and Industrial Research, India (MLP-2005) and ILS, Bhubaneswar intramural grant.

## ACKNOWLEDGEMENTS

In memory of all who have lost their lives to COVID-19. We gratefully acknowledge the authors from the originating laboratories of the submitted samples in NCBI SRA and also the COVID-19 teams of CSIR-IGIB Delhi, ILS Bhubaneswar, CSIR-CCMB and NCDC for their efforts in aggregating, processing and sharing transcriptomic data of COVID-19 infected individuals. We also acknowledge the authors from the originating laboratories of the submitted SARS-CoV-2 genome data in GISAID. We would also like to thank Bharati Singh and Gulam Hussain Syed for providing cultured Vero E6 cells, Khushboo Singhal for her inputs and Pragyan Acharya and Abhay Sharma for their critical reviews of the manuscript.

## AUTHOR CONTRIBUTIONS

Conceptualization: MM, SKR; Pipeline Design: AKP, TA, BU; Phase 1 Data Processing and Analysis: AKP, GPM; Visualization: AKP, SF^1^, MM; Spike Hyper-editing Analysis: SF^1^, LT, AKP; North Indian Cohort Sample Curation: BU, MF; East Indian Cohort Sample Curation and Processing: GPM, SW, AG, SKR; South Indian Cohort Sample Curation and Processing: SB, DTS; Phase 2 Sample Curation and Processing: GPM, SW, AK, AJ, SF^2^, SA, BU, MSD, RM, RVS, KP, SK, PR; Cultured Vero Cell Processing and qRT-PCR: AJ, GPM; Replicate Sample Curation: RCB, AJ, MKD, MI; Writing (Original Draft): MM, AKP, LT, SF^1^, GPM; Writing (Review and Editing): LT, SKR, DTS, MF, MM, AKP, GPM.

## COMPETING INTERESTS

Authors declare no conflict of interest.

## SUPPLEMENTARY DATA

**Supplementary Figure S1.**
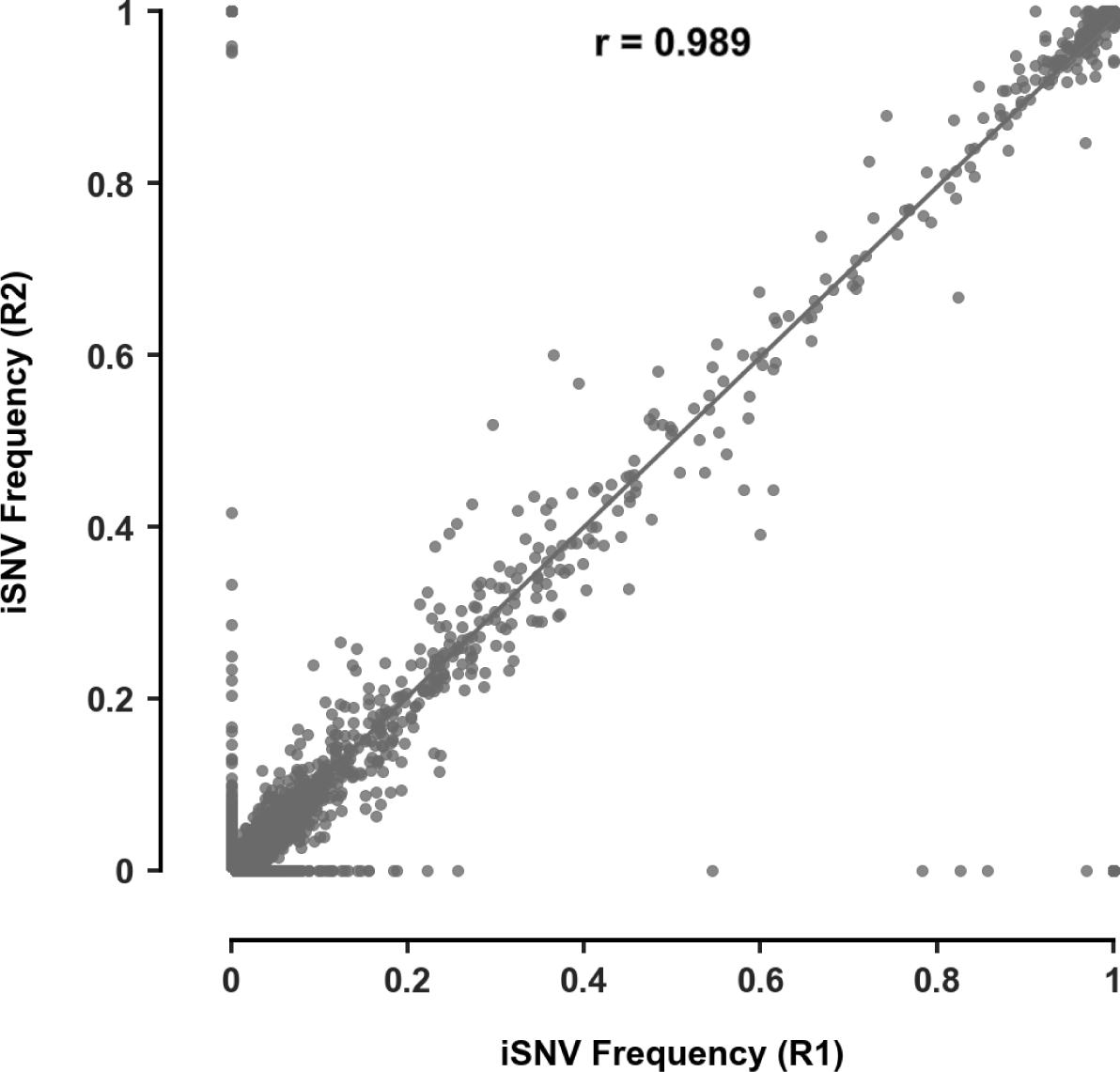
iSNV concordance between replicates. Correlation plot illustrating the concordance in iSNV frequencies between replicates (n=500).

**Supplementary Figure S2.**
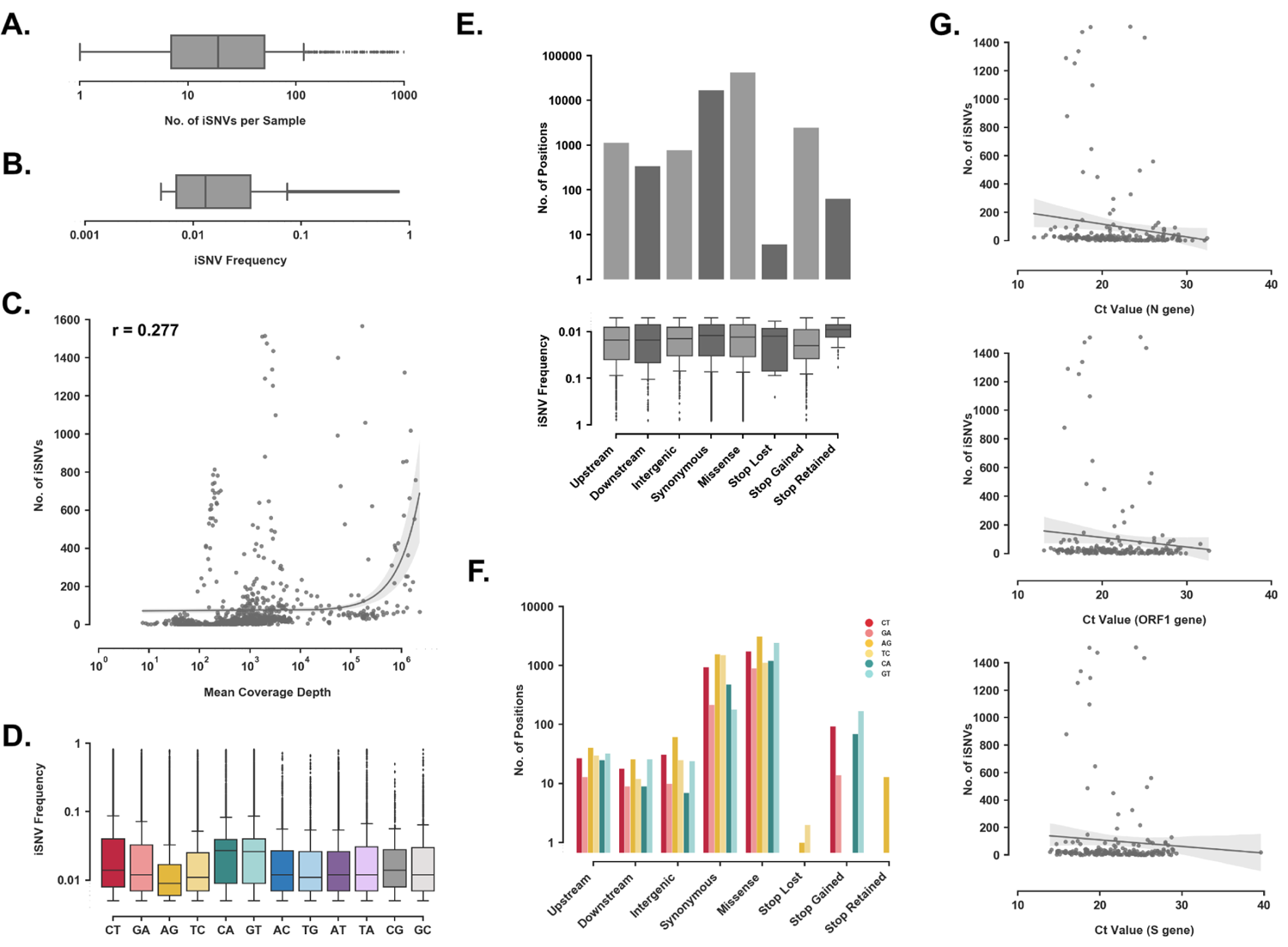
Distribution of iSNVs in Phase 1 samples. (**A**) Distribution of the number of iSNVs per sample (n=945) (**B**) Distribution of the frequencies of all captured iSNVs (n=79068) (**C**) Scatter plot to assess the correlation between number of iSNVs and mean coverage depth in samples (n=945) (**D**) Distribution of iSNV frequencies with respect to the nucleotide change (**E**) Split plot showing distribution of number of iSNVs and its potential impact vis-a-vis nature of amino acid substitution (n=62971) (**F**) Distribution of nucleotide change mediated by iSNVs with respect to nature of amino acid sequence change (**G**) Correlation plot to compare the concordance between the number of iSNVs in samples and Ct values of N, ORF1 and S gene respectively.

**Supplementary Figure S3.**
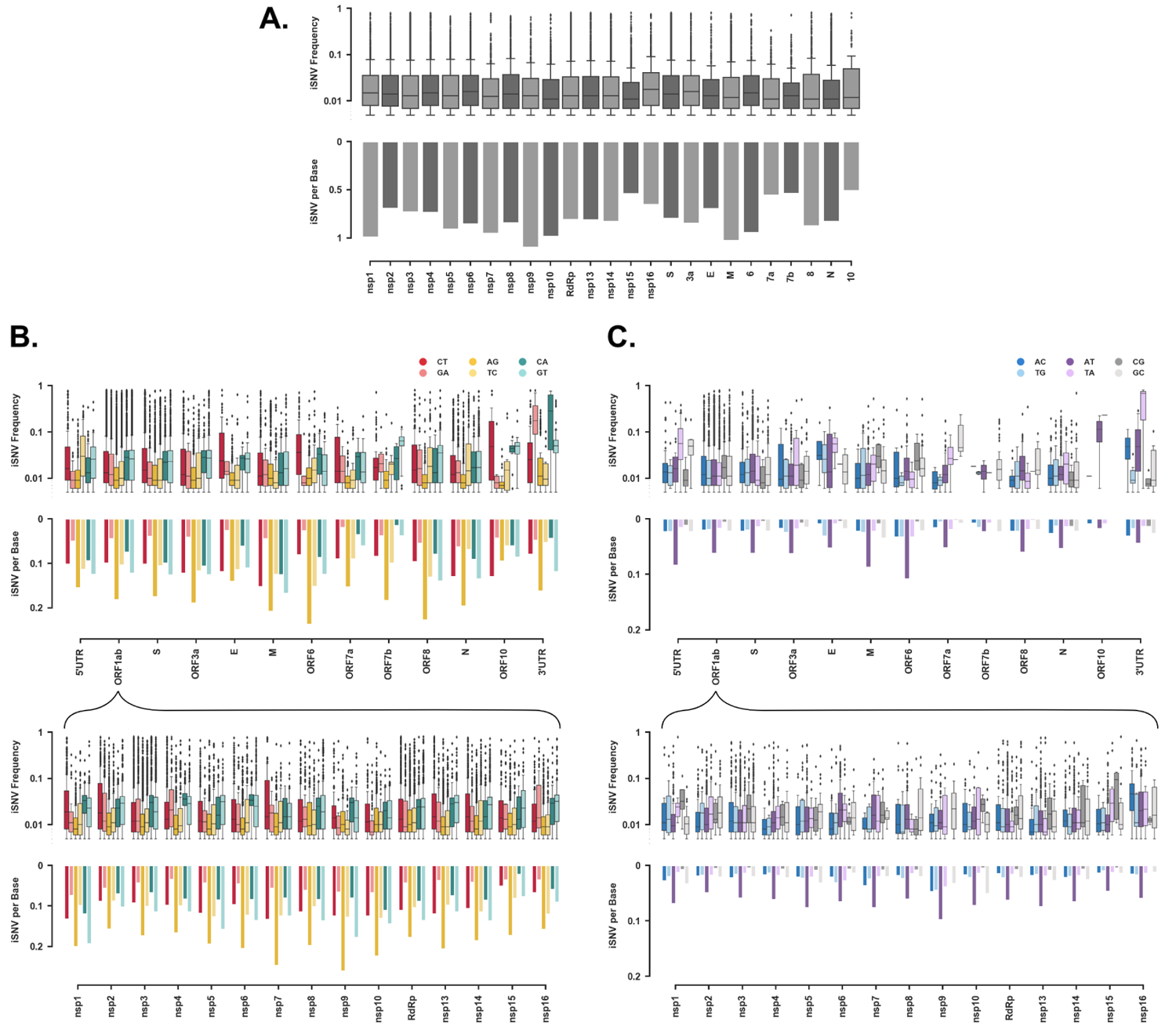
Distribution of iSNVs across the SARS-CoV-2 genome. (**A**) Split plot depicting the distribution of iSNV frequencies and number of iSNVs per base in each protein-coding domain. (**B**) Split plot depicting the distribution of iSNV frequencies and number of iSNVs per base in each protein-coding domain with respect to potential APOBEC (CT/GA), ADAR (AG/TC) and ROS (CA/GT) inflicted nucleotide changes. (**C**) Split plot depicting the distribution of iSNV frequencies and number of iSNVs per base in each protein-coding domain with respect to other nucleotide changes.

**Supplementary Figure S4.**
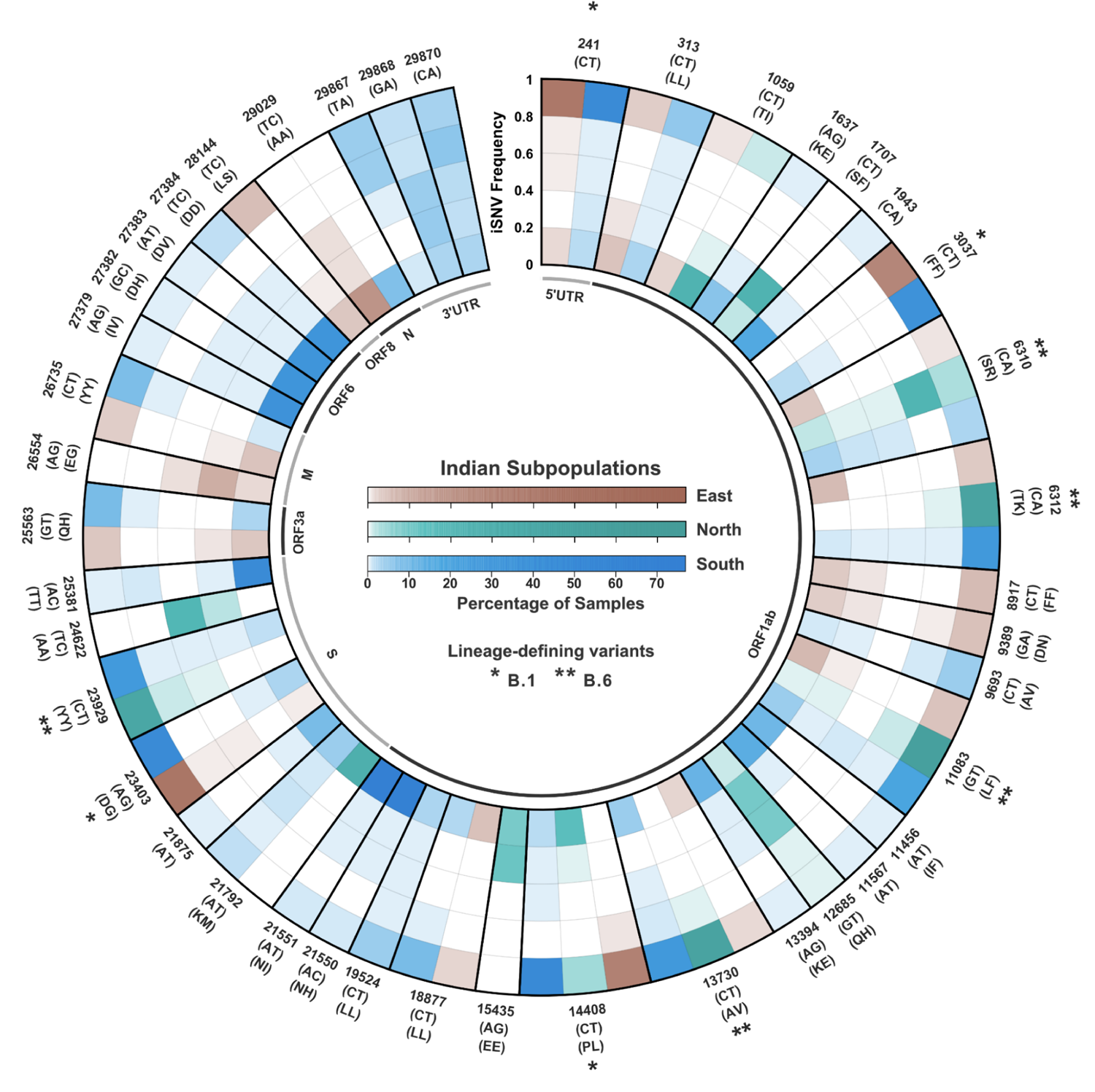
Spectrum of iSNVs in Indian subpopulations. Radial plot showcasing frequency distribution for select iSNV sites in East India, North India and South India denoted in Brown, Green and Blue respectively. Each concentric ring depicts an iSNV frequency range of 0.2 and the different colour gradients depict different populations with percentage of samples at a given position in each bin. The outer labels denote the position of change, nucleotide change and amino acid change. Variations that define the B.1 and B.6 lineages have been marked (*) and (**) respectively.

**Supplementary Figure S5.**
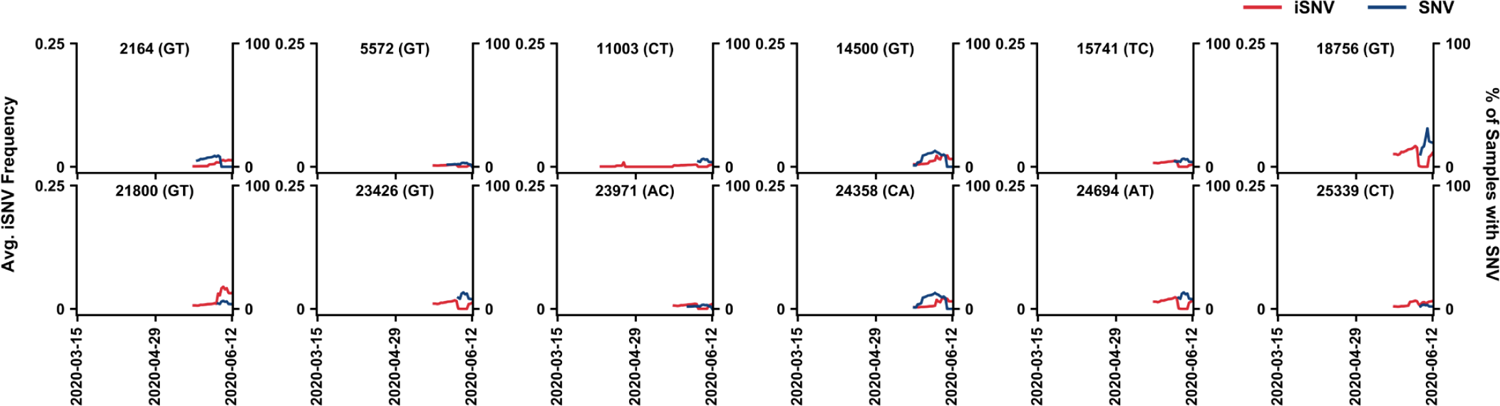
Spatio-temporal dynamics of iSNVs in East India. Line plot showing temporal trends of iSNV frequency and incidence of SNV in the East India cohort. The left y-axis denotes the average iSNV frequency and the right y-axis denotes the percentage of samples with SNVs on a 14-day rolling basis. Red and blue lines illustrate the average iSNV frequency and the percentage of samples with SNVs at the site respectively.

**Supplementary Figure S6.**
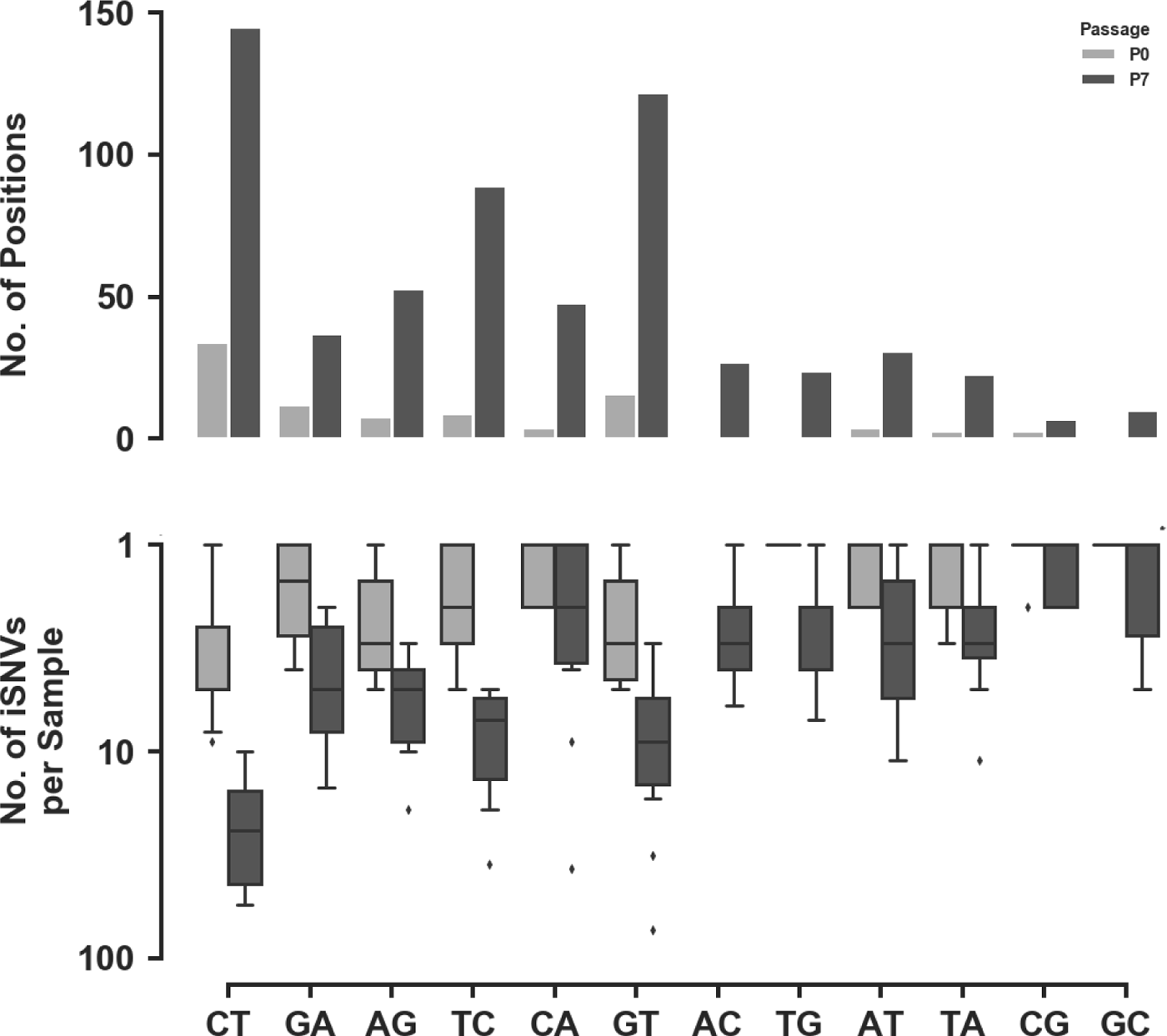
Accumulation of iSNVs in Vero cells. Comparison of iSNV prevalence at the 0th and the 7th passage of Vero cells infected with independent isolates of SARS-CoV-2 from 11 samples in the East India cohort.

**Supplementary Table S1:** **Type of amino acid change at Spike iSNV sites in hyper-edited samples.** The iSNV associated Spike variants in the hyper-edited samples characterised according to hydrophobic, polar and charged (positive and negative) nature of the residue. The rows indicate the amino acid property of the original residue and the column indicates the property of the modified residue. The number of iSNV associated variants falling in each of the pair categories are listed.

|  | Hydrophobic | Polar | Positive | Negative |
| --- | --- | --- | --- | --- |
| Hydrophobic | 45 | 31 | 3 | 4 |
| Polar | 14 | 19 | 21 | 6 |
| Positive | 11 | 2 | 7 | 2 |
| Negative | 19 | 6 | 0 | 2 |

## Supplementary Dataset Legends

**Supplementary Dataset S1:** Coordinates of homopolymeric regions in the SARS-CoV-2 genome

**Supplementary Dataset S2:** Data summary of pipeline analyses and metadata for each sample

**Supplementary Dataset S3a:** Phase 1 sample-wise frequencies of all recorded iSNVs in global populations

**Supplementary Dataset S3b:** Phase 1 sample-wise frequencies of all recorded iSNVs in Indian subpopulations

**Supplementary Dataset S4:** Summary of nucleotide change, codon change and amino acid sequence change of all recorded iSNV sites in Phase 1 samples and the number of samples with respective SNVs in GISAID

**Supplementary Dataset S5:** qRT-PCR results for mRNA expression of APOBEC3B and ADARB1 genes

**Supplementary Dataset S6:** Sample-wise frequencies of all recorded iSNVs in SARS-CoV-2 infected cultures of Vero E6 cells

**Supplementary Dataset S7:** iSNVs mapped to lineage-defining variants of VOCs and VOIs listed by the WHO by 30th June 2021

**Supplementary Dataset S8:** iSNVs mapped to potential immune-escape variants

## Notes

### Competing Interest Statement

The authors have declared no competing interest.

